# Can floral nectars reduce transmission of *Leishmania?*

**DOI:** 10.1101/2021.10.08.463579

**Authors:** Evan C Palmer-Young, Ryan S Schwarz, Yanping Chen, Jay D Evans

## Abstract

Insect-vectored *Leishmania* are the second-most debilitating of human parasites worldwide. Elucidation of the environmental factors that affect parasite transmission by vectors is essential to develop sustainable methods of parasite control that do not have off-target effects on beneficial insects or environmental health. Many phytochemicals that inhibit growth of sand fly-vectored *Leishmania*—which have been exhaustively studied in the search for phytochemical-based drugs—are abundant in nectar, which provide sugar-based meals to infected sand flies. In a quantitative meta-analysis, we compare inhibitory phytochemical concentrations for *Leishmania* to concentrations present in floral nectar and pollen. We show that nectar concentrations of several flowering plant species exceed those that inhibit growth of *Leishmania* cell cultures, suggesting an unexplored, landscape ecology-based approach to reduce *Leishmania* transmission. Strategic planting of antiparasitic phytochemical-rich floral resources or phytochemically enriched baits could reduce *Leishmania* loads in vectors, providing an environmentally friendly complement to existing means of disease control.

## INTRODUCTION

Plant secondary metabolites have a long history of use against human disease and provide the basis for both traditional medicines and many modern drugs (1), including treatments for neglected tropical diseases (2). The sand fly-vectored *Leishmania* parasites cause disease in >2 million humans each year, with 10% of the world’s population at risk, and have a greater health burden than any human parasite besides malaria (3). These Infections include >0.2M infections with visceral leishmaniasis, which, if untreated, results in >90% patient mortality (3, 4). Due to their clinical significance, *Leishmania* spp. have been studied intensively in a search for affordable and effective treatments for human infections (5), including exhaustive testing of plant extracts and their components against both mammal- and insect-associated parasite life stages (2, 6). These studies have suggested new treatments for trypanosomatid-associated infections of humans (7) and related parasites of beneficial insects (8, 9).

As in humans, antimicrobial phytochemicals can enhance resistance to infection in plants themselves (10) and in other plant-consuming animals, including insects (11). The diets of blood-feeding, disease-vectoring insects such as sand flies and mosquitoes include plant sugars as well as blood (12).

Different plant sugar sources can profoundly affect the development of *Plasmodium falciparum* malaria in *Anopheles* mosquitoes (13) and *Leishmania major* in sand flies (*Phlebotomus papatasi*) (14). Sand flies feed on plant sugars between acquisition and transmission of *Leishmania* to humans and other mammals (15). Although sand flies may acquire these meals from plant sap, fruit, or aphid- or cicada-derived honeydew, and fruits (16), floral nectar appears to be a preferred food source (12).

The role of nectar chemistry in insect disease ecology has recently been highlighted by work on infections of pollinators. Floral nectar and pollen, their constituent secondary metabolites, and the composition of flowering plant communities can ameliorate trypanosomatid growth and infection in bumble bees (8, 9, 17–19). Both nectar and pollen—which may mix with and influence the chemistry of nectar at flowers (20)—contain diverse secondary metabolites that shape plant-pollinator ecology and plant-microbe ecology (21–25). Flavonoids are one class of antimicrobial and antileishmanial compounds (26, 27) that are ubiquitous in both nectar and pollen, with concentrations in pollen often exceeding 1% of total dry matter (28, 29). This suggests that consumption of secondary metabolite-rich nectars could mitigate *Leishmania* transmission by reducing infection intensity in nectar-feeding sand fly vectors (12), pointing to a new strategy for drug- and insecticide-free disease control. However, despite appreciation for the clinical antileishmanial potential of plant metabolites (2), growing recognition of the role of plant metabolites—including those in nectar and pollen—in insect infection, and the critical role of plant sugars in sand fly diets, there has been surprisingly little investigation into the potential for antileishmanial phytochemicals in the diets of sand flies to mitigate *Leishmania* transmission.

To assess the potential for floral resource-associated phytochemicals to reduce vector-borne infection, we compared phytochemical concentrations previously shown to inhibit *Leishmania* to concentrations previously found in floral nectar and pollen. Our synthesis of prior work on *Leishmania* phytochemical sensitivity with nectar and pollen secondary chemistry shows that many floral nectars contain antileishmanial compounds at concentrations sufficient to inhibit parasite growth. These findings suggest an unexplored, landscape ecology-based approach to reduce transmission of widespread and virulent *Leishmania* infections. Incorporation of antiparasitic nectar sources into landscapes and domestic settings could simultaneously benefit pollinator and public health.

### ASSESSMENT OF NECTAR AND POLLEN PHYTOCHEMISTRY

We compared the flavonoid concentrations found in a previous survey of methanolic extracts from 26 floral nectars and 28 pollens (28, 30) with previously published results from *in vitro* screening of various *Leishmania* spp. (**Supplementary Table 1)**. We focused on flavonoids because these compounds were the most consistently present class of compounds across both nectar and pollen (28) and— particularly in the case of quercetin— some of the most potent and selective compounds against *Leishmania* (27, 31, 32). To prevent overestimation of inhibitory potential that could result from including flavonoids of lesser or unknown antiparasitic activity, we further distinguished between total flavonoid concentrations and those with a kaempferol, quercetin, apigenin, or luteolin aglycone, each of which has well-documented antileishmanial effects (27, 33, 34) **(Supplementary Table 1)**.

We analyzed micromolar concentrations to enable pooling across compounds with different parent flavonoids and glycosides. Although flavonoid glycosides can be less potent than their parent aglycones (27), we included flavonoid glycosides because these compounds are hydrolyzed by intestinal glucosidases—a variety of which are found in sand flies (35)—to their corresponding aglycones (36, 37). We focus our discussion on nectar because sand flies, like other Diptera, do not have chewing mouthparts that would enable direct consumption of pollen and other solid foods (38). However, incidental presence of pollen in nectar can dramatically increase the nectar’s concentrations of amino acids (39), with ecologically relevant effects on nectar-feeding insects (20).Pollen could similarly affect nectar’s phytochemical profile and antimicrobial effects. For example, presumably pollen-derived cinnamic acid-spermidine conjugates were found in nectar in pollen of two species in our previous survey—*Digitalis purpurea* and *Helianthus annuus* (28). In *H. annuus*, nectar concentrations averaged 1.7% of pollen concentrations, despite exclusion of large insects that contribute to such “contamination” (39) for 24 h prior to sampling. We therefore also discuss pollen concentrations that exceed the *Leishmania* IC50 estimates by >100-fold, on the grounds that the much (235-fold (28)) higher flavonoid concentrations found in pollen could meaningfully alter the antiparasitic activity of nectar, even when pollen is present at <1% of nectar volume.

### VULNERABILITY OF *LEISHMANIA* TO FLORAL CHEMICALS

We compiled 18 *Leishmania* IC50 estimates for 4 flavonoids—quercetin (n = 8), kaempferol (n = 4), apigenin, and luteolin (n = 3 each) that have been relatively well studied for effects on *Leishmania* spp. cell cultures **(Supplementary Table 1)**. Most (11 of 18) of the assays used the promastigote (i.e., insect-associated) life stage; the remainder used either intracellular (n = 4) or axenic (n = 3) amastigotes **(Supplementary Table 1)**. These *Leishmania* IC50 estimates were then compared to the flavonoid concentrations found in a previous survey of secondary metabolites of nectar and pollen (28).

Flavonoids were found in the nectar of 21 of 26 species (81%) and in pollen of 26 of 28 species (93%), accounting for 30% of the total phytochemical content in nectar and 41% in pollen (28). Total flavonoid concentrations exceeded 100 μM in nectar from 8 of 26 nectars (31%, median concentration 30.9 μM, IQR 4.24-127 μM; median 61.4 μM after exclusion of the five species without nectar flavonoids) and exceeded 10^4^ μM in 18 of 28 pollens (64%, median 1.36 · 10^4^ μM, IQR 4.31 · 10^3^ to 2.30 · 10^4^ μM) **(Figure 1)**. Glycosides of quercetin (found in 9 of 26 nectars and 14 of 28 pollens) and kaempferol (5 of 26 nectars and 19 of 28 pollens) were most common (28).

**Figure 1.**
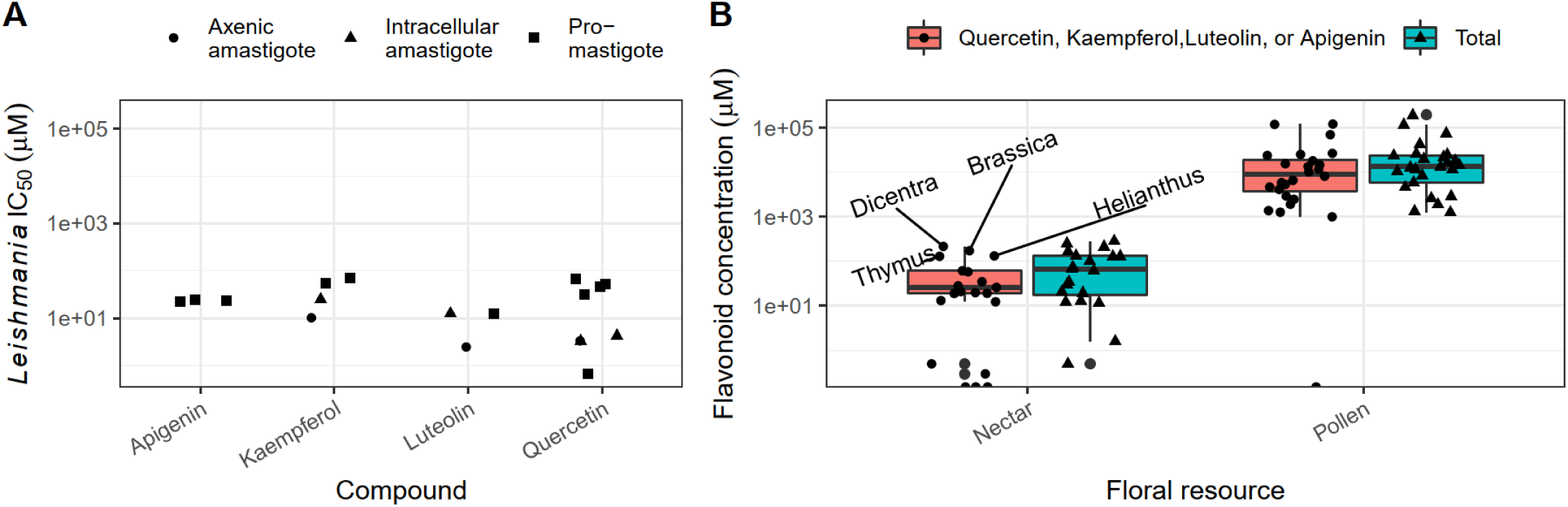
Published *Leishmania* IC50 estimates for selected flavonoids (A) relative to concentrations of the corresponding compounds in nectar and pollen (B). Shapes in panel (A) correspond to the *Leishmania* stage tested. Boxplots in panel (B) show medians and interquartile ranges for concentrations of quercetin, kaempferol, apigenin and luteolin derivatives (red boxes) total flavonoids (blue boxes). Points show median concentrations (pooled across individual samples) by species. Text annotations denote species with >100 µM of the selected flavonoids in nectar (*Brassica napus, Dicentra eximia, Helianthus annuus*, and *Thymus vulgaris*). Literature references for *Leishmania* IC50 estimates are given in **Supplementary Table 1**.

Compounds with a parent aglycone of quercetin, kaempferol, apigenin, or luteolin accounted for 62% of flavonoid compounds and 54% of molar concentrations in nectar, and 72% of compounds and 75% of molar concentrations in pollen. In nectar, median concentration of this subset of compounds across all species (20.3 μM, IQR 9.37-58.9 μM) was remarkably close to the 23.1 μM median IC50 for *Leishmania* (based on 18 references **(Supplementary Table 1)**). Concentrations exceeded 100 μM (i.e., more than the highest *Leishmania* IC50 for any of the parent compounds) in nectar from 4 of 26 species (*Dicentra eximia, Brassica napus, Helianthus annuus, and Thymus vulgaris*). In pollen, median concentrations exceeded 10^4^ μM (i.e., >100-fold the greatest *Leishmania* IC50) in pollen from 12 of 28 species, including two species (*Lythrum salicaria* (1.21 · 10^5^) and *Solidago canadensis* (1.19 · 10^5^)) with concentrations >10^5^ μM—over three orders of magnitude above the greatest *Leishmania* IC50 **(Figure 1**).

Antileishmanial compounds in nectar were not limited to flavonoids. Seven nectars contained chlorogenic acid, with a median concentration (51.2 μM) similar to the IC50 for *L. donovani* promastigotes (54 μM (40)) and 100-fold greater than the IC50 for *L. amazonensis* promastigotes (0.5 μM (41)). The species with the highest median concentration of chlorogenic acid (*Dicentra eximia*, 184 μM) also had the highest concentration of the selected flavonoids **(Figure 1)**. Nectar concentrations of two additional species (*Penstemon digitalis*, 134 μM) and *Rhododendron prinophyllum* (56.7 μM) also exceeded the *L. donovani* promastigote IC50 **(Figure 2)**. Chlorogenic acid was also found in seven pollens at up to 3760 μM (*Persea americana*), with a median concentration (1227 μM) over 20-fold greater than that found in nectar and over three orders of magnitude above the *L. amazonensis* promastigote IC50 (41) **(Figure 2)**.

**Figure 2.**
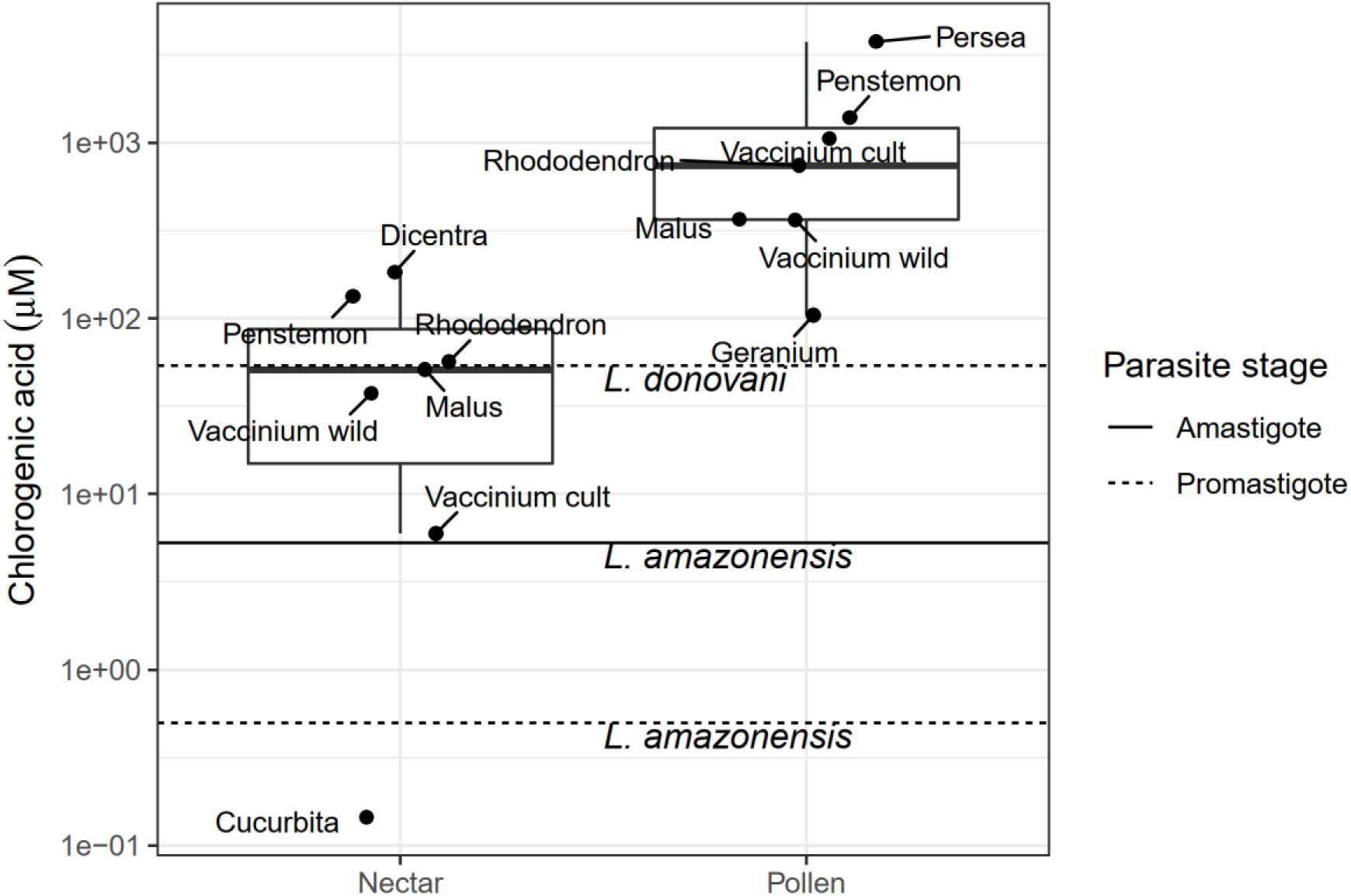
Concentrations of chlorogenic acid in nectar and pollen in comparison with inhibitory concentrations for *Leishmania*. Points represent median concentrations from species with detectable chlorogenic acid (sampled in (28)). Horizontal lines show published IC50 values (40, 41). Sampled plant species (labeled by genus) were *Dicentra eximia, Penstemon digitalis, Rhododendron prinophyllum, Malus domestica, Vaccinium corymbosum, Cucurbita pepo, Persea americana, and Geranium maculatum*. For *Vaccinium*, “cult” refers to cultivars and “wild” refers to wild plants.

Nectar of one species (*Thymus vulgaris*) contained the caffeic acid-dihydroxyphenyl lactic acid ester rosmarinic acid. Median concentration (165 μM, IQR 87.7 - 206 μM) was 10-fold greater than the IC50 for *L. donovani* promastigotes (16.3 μM (40))—against which rosmarinic acid and apigenin were the most selective of the compounds evaluated—over 30-fold greater than the 4.8 μM IC50 for *L. amazonensis* amastigotes (41), and over 200-fold greater than the 0.7 μM reported for *L. amazonensis* promastigotes (41) **(Figure 3)**. Nectar of *T. vulgaris* is also notable for its high thymol content (26.1 μg mL^-1^ (42)), which exceeds six of the eight IC50 values reported for *Leishmania* promastigotes (**Supplementary Table 1**, (43, 44)).

**Figure 3.**
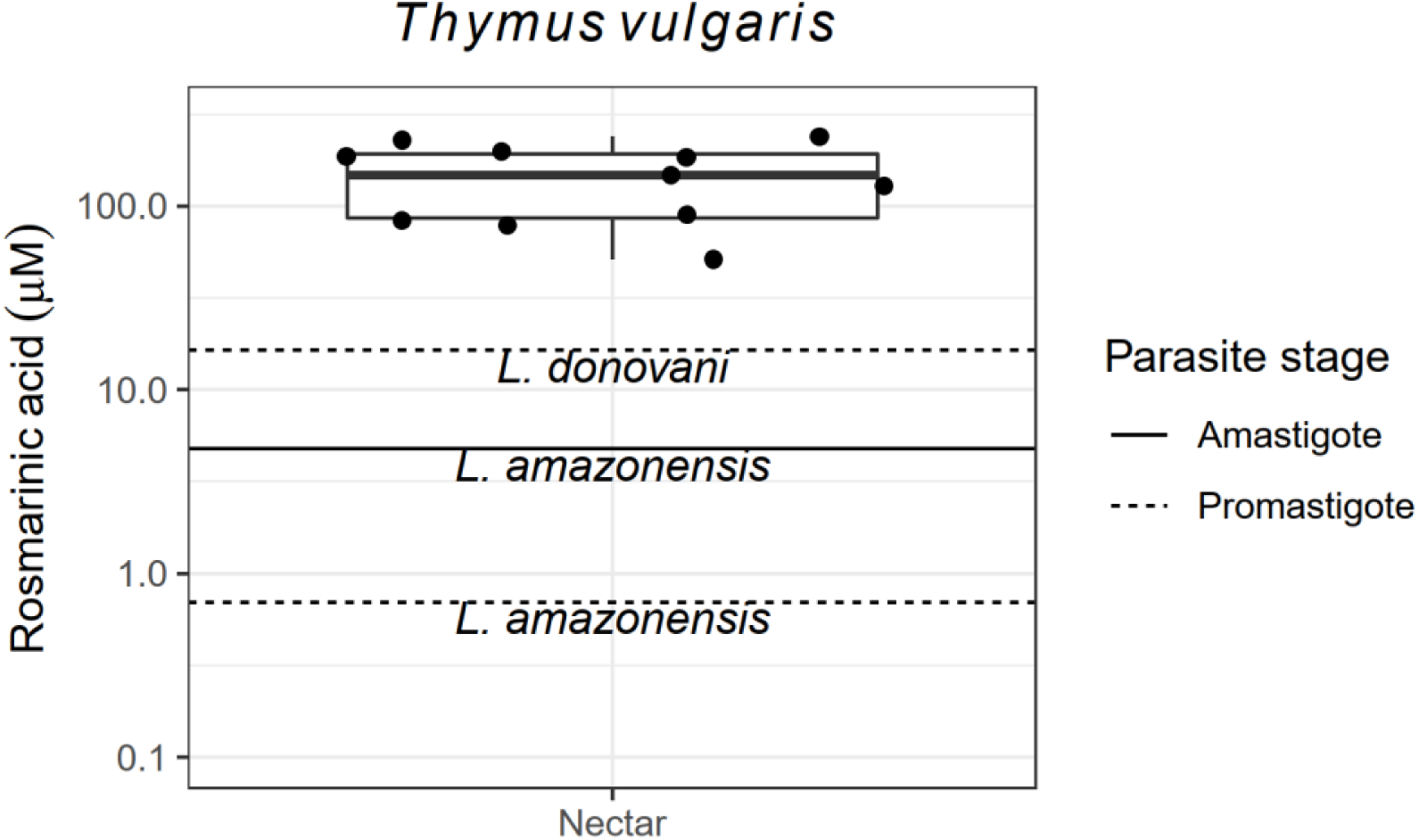
Concentrations of rosmarinic acid in *Thymus vulgaris* nectar in comparison with inhibitory concentrations for *Leishmania*. Points represent individual nectar samples (from (28)). Horizontal lines show published IC50 values (40, 41).

## APPLICATIONS

Our synthesis of a previous survey on the quantitative phytochemical composition of nectar and pollen with the extensive body of research on phytochemical-mediated inhibition of Leishmania *in vitro* reveals the potential for floral resources to ameliorate vector-mediated transmission of *Leishmania*, the second most disabling human parasite worldwide (3). The most common compounds in nectar and pollen—flavonoids and their glycosides—have shown strong inhibitory effects against *Leishmania* (27, 32, 45). Our findings indicate that a subset of the floral nectars analyzed to date—including the common garden herb *Thymus vulgaris* (thyme) and the widespread crop species *Helianthus annuus* (cultivated sunflower) contain bioactive flavonoids at concentrations that inhibit growth of diverse *Leishmania in vitro*. Incidentally, both plant species have also been shown to mitigate transmission and infectivity of bumble bee trypanosomatids, including in field mesocosms and landscape surveys (18, 19). Deliberate encouragement of these and other plants with trypanosomatid-inhibiting chemistry could reduce infection intensity in disease-vectoring hematophagous insects as well.

The effect of sugar-containing meals from plant sources was a long-overlooked component *of* sand fly ecology that proved crucial in *Leishmania* transmission (15). However, feeding of sand flies on several plant taxa causes marked parasite mortality—up to 88% in the case of castor bean (*Ricinus communis*) (14), the lectins of which agglutinate a variety of insect trypanosomatids (46)—paralleling the strong effects of plant sugar sources on malaria infection in mosquitoes (13). Although flowering plant nectar sources might at first glance appear to be a liability for *Leishmania* transmission due to the food they provide to sand flies, sugar starvation in fact results in greater vector infection intensity and natural selection for flies with lesser parasite resistance (47, 48). This finding is consistent with the preponderance of *Leishmania* hotspots in arid regions, where plant sugar sources are scarce (47, 48).

High parasite loads also alter sand fly feeding on mammals in ways that promote transmission to new hosts (49). These lines of evidence suggest that despite their role as vector food sources, phytochemical-rich floral nectar sources could have a net transmission-reducing effect. Given the apparent susceptibility *of Leishmania* to common nectar compounds at ecologically relevant concentrations, the high *Leishmania* infection intensity in flies from regions lacking in sugar sources (47), and the numerous nutritional benefits of floral resources and constituent phytochemicals for bees (50), landscapes containing plants rich in trypanosomatid-inhibiting nectars could mutually benefit pollinator and public health.

Although currently a matter of speculation, the presence of pollinators could conceivably enhance the antiparasitic properties of nectar for disease-vectoring nectar consumers by introducing compounds from pollen or floral tissues (51, 52)—a potential ecosystem service that is currently entirely unexplored. Introduction of pollen as a result of bee visitation can increase nectar amino acid concentrations by an order of magnitude, and potentially introduce additional compounds from plants of other species as well (39). Given that flavonoid concentrations in pollen are 200-fold higher than those in nectar (28), incidental ingestion of even small amounts of pollen could substantively inhibit proliferation of parasites and the transmission potential of their vectors. In *H. annuus*, pollen-associated spermidines occurred at concentrations >1% of those in pollen even when pollinators were excluded (28). In our meta-analysis, eight of the 28 pollens previously surveyed contained flavonoids at concentrations that exceeded 100-fold the maximum inhibitory concentration reported for *Leishmania* **(Figure 2)**. This suggests that as little as 1% incidental addition of pollen to nectaries would be sufficient for *Leishmania* inhibition, even for nectars that lack antileishmanial flavonoids initially. Such phytochemically enriched nectars could have particular potency against *Leishmania*, which appear both sensitive to flavonoids and, given their establishment in the midgut and forward migration in the alimentary canal (15), directly exposed to ingested compounds before appreciable metabolism of these compounds by hosts or their hindgut microbiota. The limited intestinal absorption of ingested flavonoids (37), hydrolysis of glycosides found in plants to their more potent aglycones in the intestine (35, 37), and likelihood of direct contact between parasites in the anterior midgut and ingested phytochemicals all indicate the potential for flavonoid-rich nectars to reduce *Leishmania* infection in sand flies.

## CONCLUSIONS

The global toll of *Leishmania* infection and the difficulties of eradicating its sand fly vectors and non-human reservoirs demand the development of new, environmentally compatible strategies to reduce parasite transmission (53). Reduction of transmission *via* supply of antiparasitic nectar sources in local landscapes could have profound significance for public health—reducing reliance on drug treatments that may be costly, inaccessible, or potentially hazardous (3)—while simultaneously supporting populations of beneficial insects and their resistance to insect-specific trypanosomatid infections (17, 18). The fields of insect ecology and medicinal chemistry for insect-vectored parasites have thus far developed more in parallel than in concert. Integrating knowledge of medicinal plant chemistry and plant-mediated tritrophic interactions that affect parasites in disease-vectoring insects holds promise for environmentally friendly control of trypanosomatid threats to global health.

## ACKNOWLEDGMENTS

This project was supported by the USDA Agricultural Research Service Beltsville Bee Research Laboratory in house fund; USDA-NIFA Pollinator Health Grant 2020-67013-31861 to JDE and YPC; and a North American Pollinator Protection Campaign Honey Bee Health Improvement Project Grant and an Eva Crane Trust Grant to ECPY and JDE. Funders had no role in study design, data collection and interpretation, or publication. We thank the anonymous reviewers for their service in improving the manuscript.

## DATA AVAILABILITY

All data are supplied in the Supplementary Information, Data S1.

## AUTHORS’ CONTRIBUTIONS

ECPY, JDE, and RSS conceived the study. ECPY conducted experiments, analyzed data, and drafted the manuscript with guidance from JDE, YC, and RSS. All authors revised the manuscript and gave approval for publication.

