## Supplementary information for "Can floral nectars reduce transmission of *Leishmania?*"

Running title: Nectar phytochemicals and *Leishmania* transmission

Evan C Palmer-Young [1]*, Ryan S Schwarz [2], Jay D Evans [1]

[1] USDA-ARS Bee Research Lab, Beltsville, MD, USA

[2] Department of Biology, Fort Lewis College, Durango, CO, USA

### SUPPLEMENTARY TABLES

**Supplementary Table 1.** Reported inhibitory concentrations (IC50) for the flavonoids apigenin, luteolin, kaempferol, and quercetin; and the esters chlorogenic and rosmarinic acids. Species are listed as in the original publications, but note that *L. chagasi* is considered synonymous with *L. infantum* (1).

| Substance | Species | Stage | IC50 (µg/mL) | Assay | Duration (h) | Reference | Notes |
|---|---|---|---|---|---|---|---|
| Apigenin | *Leishmania amazonensis* | P | 6.4 | Cell counts | 24 | (2) | |
| Apigenin | *Leishmania mexicana* | P | 6.6 | Resazurin fluorescence | 72 | (3) | |
| Apigenin | *Leishmania donovani* | P | 6.1 | MTT absorbance | 72 | (4) | |
| Chlorogenic acid | *Leishmania amazonensis* | P | 0.2 | MTT absorbance | 72 | (5) | |
| Chlorogenic acid | *Leishmania amazonensis* | IA | 1.9 | Cell counts | 48 | (5) | |
| Chlorogenic acid | *Leishmania donovani* | P | 19.1 | MTT absorbance | 72 | (4) | |
| Kaempferol | *Leishmania donovani* | AA | 2.9 | Resazurin fluorescence | 72 | (6) | |
| Kaempferol | *Leishmania donovani* | IA | 7.15 | Cell counts | 72 | (7) | |
| Kaempferol | *Leishmania peruviana* | P | 20.4 | Cell counts | 72 | (8) | |
| Kaempferol | *Leishmania braziliensis* | P | 15.3 | Cell counts | 72 | (8) | |
| Luteolin | *Leishmania donovani* | P | 3.6 | Cell counts | 24 | (9) | |
| Luteolin | *Leishmania donovani* | IA | 3.6 | Cell counts | 24 | (9) | |
| Luteolin | *Leishmania donovani* | AA | 0.7 | Resazurin fluorescence | 72 | (6) | |
| Quercetin | *Leishmania amazonensis* | P | 9.4 | Cell counts | 48 | (10) | |
| Quercetin | *Leishmania amazonensis* | P | 0.2 | MTT absorbance | 72 | (5) | |

| | | | | | | |
|---|---|---|---|---|---|---|
| Quercetin | *Leishmania amazonensis* | IA | 1.3 | Cell counts | 48 | (5) |
| Quercetin | *Leishmania amazonensis* | IA | 1 | Cell counts | 72 | (11) |
| Quercetin | *Leishmania donovani* | P | 13.7 | Cell counts | 24 | (9) |
| Quercetin | *Leishmania donovani* | AA | 1 | Resazurin fluorescence | 72 | (6) |
| Quercetin | *Leishmania peruviana* | P | 16.1 | Cell counts | 72 | (8) |
| Quercetin | *Leishmania braziliensis* | P | 20.7 | Cell counts | 72 | (8) |
| Rosmarinic acid | *Leishmania amazonensis* | P | 0.2 | MTT absorbance | 72 | (5) |
| Rosmarinic acid | *Leishmania amazonensis* | IA | 1.7 | Cell counts | 48 | (5) |
| Rosmarinic acid | *Leishmania donovani* | P | 5.9 | MTT absorbance | 72 | (4) |
| Thymol | *Leishmania amazonensis* | P | 26.8 | Resazurin fluorescence | 24 | (12) |
| Thymol | *Leishmania infantum* | P | 12.85 | Bio-luminescence | 24 | (13) |
| Thymol | *Leishmania infantum* | IA | 23.93 | ELISA | 48 | (13) |
| Thymol | *Leishmania chagasi* | P | 9.8 | MTT absorbance | 72 | (14) |
| Thymol | *Leishmania chagasi* | P | 9.8 | MTT absorbance | 72 | (15) |
| Thymol | *Leishmania infantum* | P | 7.2 | MTT absorbance | 24 | (16) |
| Thymol | *Leishmania amazonensis* | P | 19.5 | Cell counts | 48 | (17) |
| Thymol | *Leishmania chagasi* | P | 65.2 | Cell counts | 72 | (18) |

Abbreviations:

P: Promastigote

AA: Axenic amastigote

IA: Intracellular amastigote

MTS: [3-(4, 5 dimethyl-thiazol-2-yl) 5- (3-carboxymethoxyphenyl)-2-(4-sulphonyl)-2H-tetrazolium]"

PMS: phenazine methosulfate

MTT: 3-[4,5-dimethylthiazol-2-yl]-2,5-diphenyltetrazolium bromide

### SUPPLEMENTARY DATA

**Zipped folder with data spreadsheets** for *Leishmania* inhibitory concentrations (**leishmania_ic50**) and nectar and pollen flavonoid concentrations (**nectar.pollen.flavonoids**).
